# Autophagy limits inflammatory gene expression through targeting of nuclear p65/RelA by LC3 and p62 for lysosomal degradation

**DOI:** 10.1101/2022.11.02.514846

**Authors:** Cristina Brischetto, Patrick Mucka, Eva Kaergel, Claus Scheidereit

**Affiliations:** Signal Transduction in Tumor Cells, Max-Delbrück-Center for Molecular Medicine, Robert-Rössle-Str. 10, 13092 Berlin, Germany

**Keywords:** autophagy, lysosome, proteasome, chronic inflammation, NF-κB, transcription, degradation

## Abstract

The interplay between NF-κB signaling and autophagy regulates inflammatory signaling in different cellular contexts and in response to different stimuli. The impairment of this crosstalk may play a role in chronic inflammation and in tumorigenesis. However, the molecular mechanism by which these two pathways interact to regulate the inflammatory response remains elusive. By using biochemical analysis and imaging techniques, we characterized the interaction of the endogenous autophagic marker LC3 and NF-κB/p65 in response to different stress conditions. Following irradiation or TNFα stimulation, nuclear accumulation of LC3 strongly co-localized with p65, suggesting that nuclear p65 is targeted for autophagic degradation. Mechanistically, we showed that the nuclear p65-LC3 interaction is mediated by ubiquitination of the same p65, which is recognized by the cargo receptor p62, resulting in its cytoplasmic export and lysosomal proteolysis. Accordingly, autophagy inhibition by depletion of the essential autophagy gene *ATG16L1* selectively stabilizes nuclear p65, in turn enhancing NF-κB gene expression of pro-inflammatory cytokine. Our results revealed a novel molecular mechanism that modulates the NF-κB inflammatory response through nuclear sequestration of the NF-κB/p65 subunit by autophagy proteins. These findings are of importance for developing novel therapeutic strategies against chronic inflammatory diseases displaying defective autophagy and constitutive NF-κB activity.

## Introduction

Tight regulation of immune and stress responses is essential for maintaining cellular homeostasis and resolving inflammation. Malfunction of a well-coordinated immune response can lead to chronic inflammation which, along with infection, is estimated to represent approximately a quarter of all cancer-causing factors ^1^. Activation of the transcription nuclear factor (NF)-κB plays a central role in coordinating immune responses through the induction of numerous genes, including pro-inflammatory cytokines, chemokines, proteases, and inducible nitric oxide (iNOS) ^2^. In line with its physiological importance, aberrant NF-κB activity is implicated in many pathological conditions including chronic inflammatory diseases and cancer ^3,4^. When inactive, NF-κB proteins are held in the cytoplasm by associating with inhibitory members of the IκB-family ^5^. In order to translocate into the nucleus and exert their transcriptional functions, NF-κB proteins need to be released from IκBα in a stimulus-dependent manner. A variety of stimuli, such as inflammatory cytokines, radiation, and certain pathogens lead to rapid activation of NF–κB RelA/p65-p50, the most prevalent heterodimer ^6,7^. This is triggered by the upstream IκB kinase (IKK) complex that phosphorylates IĸBα, leading to its ubiquitination and proteasomal degradation ^8^. In addition to the well-documented NF-κB–regulated resynthesis of IκBα, ubiquitin-mediated proteasomal degradation of the NF-κB subunit p65 has been proposed to force prompt termination of the NF-κB response ^9–12^.

Autophagy is an evolutionarily conserved catabolic cellular degradation pathway involved in the maintenance of cellular homeostasis ^13^. Depending on the cellular stresses, autophagy can be either non-selective or selective. Whereas non-selective autophagy typically involves random uptake of cytoplasm, selective autophagy is responsible for specifically removing certain cellular components like protein aggregates, organelles, or pathogens ^14^. This implicates a multitude of types of autophagy that play an important role in different catabolic and anabolic cellular processes. Indeed, defects in the autophagy process are responsible for many human diseases ^15,16^. Interestingly, genome-wide association studies suggested a connection between autophagy-related gene polymorphisms and many inflammatory disorders, in which NF-κB is constitutively activated ^17–19^. The interplay between autophagy and NF-κB signaling has been observed in different cellular contexts and in response to different stimuli. The impairment of this crosstalk determines cell fate and is frequently associated with tumorigenesis and tumor cell resistance to cancer therapies ^17,20–22^.

In this study, we provide a detailed molecular dissection of the functional interplay between autophagy and the NF-κB inflammatory response. We found that an interaction of LC3 with p65 in the nucleus is strongly enhanced following NF-κB activation by TNFα or γ-irradiation. The binding between p65 and LC3 depends on ubiquitin-conjugation, which is recognized by p62/SQSTM1, followed by nucleus-to-cytoplasm translocation of the p65-p62-LC3 complex and lysosomal p65 degradation. Together, we identified a novel mechanism of NF-κB response termination by autophagy proteins, providing insights that may help to improve the effectiveness of inflammation-associated cancer therapies.

## Results

### The autophagic protein LC3 localizes in the nucleus associated with NF-κB/p65 in response to stress-induced inflammation

The crosstalk between NF-κB signaling and autophagy has been suggested to regulate inflammation in different cellular contexts and in response to different stimuli ^20,21,23^. Therefore, we first investigated the effect of tumor necrosis factor-alpha (TNFα), lipopolysaccharide (LPS), and genotoxic stress, which are well-known NF-κB inducers, on the activation of autophagy in different cell types. Microtubule-associated protein light chain 3 (LC3) is an essential protein for the maturation of the autophagosome and a well-established marker for monitoring the autophagosome formation through the biochemical detection of its membrane-associated form (LC3-II) ^24^. The increase in LC3-II during and after NF-κB activation (Fig. 1a and Fig. S1a), suggested the accumulation of autophagosomes not only 1 hour after TNFα treatment but also rapidly after irradiation or LPS treatment in THP1 or MEF cells (Fig. 1a and S1a). Nuclear translocation of NF-κB subunit p65 (Fig. 1a) or the level of p65 phosphorylation (Fig. S1a) was used to monitor the activation of NF-κB. We further assessed autophagosome maturation using U2-OS cells stably expressing green fluorescent protein (GFP)-tagged LC3 and examined the presence and subcellular localization of the fluorescence signals ^24^. Quantification of LC3 puncta-like structures indicates a significant increase of autophagosome formation at 1 hour post-irradiation (IR) or post TNFα treatment compared to the untreated control and the *bona fide* induction of autophagy by nutrient starvation (EBSS), confirming TNFα- and DNA damage-induced autophagy (Fig.1b and Fig. S1b). These results were confirmed by LC3 turnover assay in the presence or not of Bafilomycin A1, an inhibitor of the late phase of autophagy, which is based on the observation that LC3-II is degraded in autolysosomes, which indicates enhancement of the autophagic flux (Fig.1c). Interestingly, we observed an accumulation of LC3-II in the nuclear fraction following stress-induced NF-κB activity (Fig.1a, b). Nuclear accumulation of LC3 has been reported to play a role in different nuclear functions in response to specific types of stress, including the turnover of nuclear factors ^25–28^. Therefore, we hypothesized that nuclear autophagy could be involved in the regulation of NF-κB upon stress-induced inflammation. To test this hypothesis, we first visualized the intracellular localization of potential interaction of LC3 with p65 by performing *in situ* proximity ligation assay (PLA) analysis, which was previously used to study the dynamics of NF-κB activation ^29^. Quantification of PLA signals revealed a significant increase of endogenous p65:LC3 binding in the nuclear compartment after stimulation with TNFα or irradiation compared to untreated cells, in which the signals remained mainly in the cytoplasm (Fig. 1d, e). Immunofluorescence analysis confirmed this finding (Fig. S1c), strongly suggesting that the NF-κB/p65 subunit and the autophagy protein LC3 interact in the nucleus after NF-κB activation.

**Fig.1:**
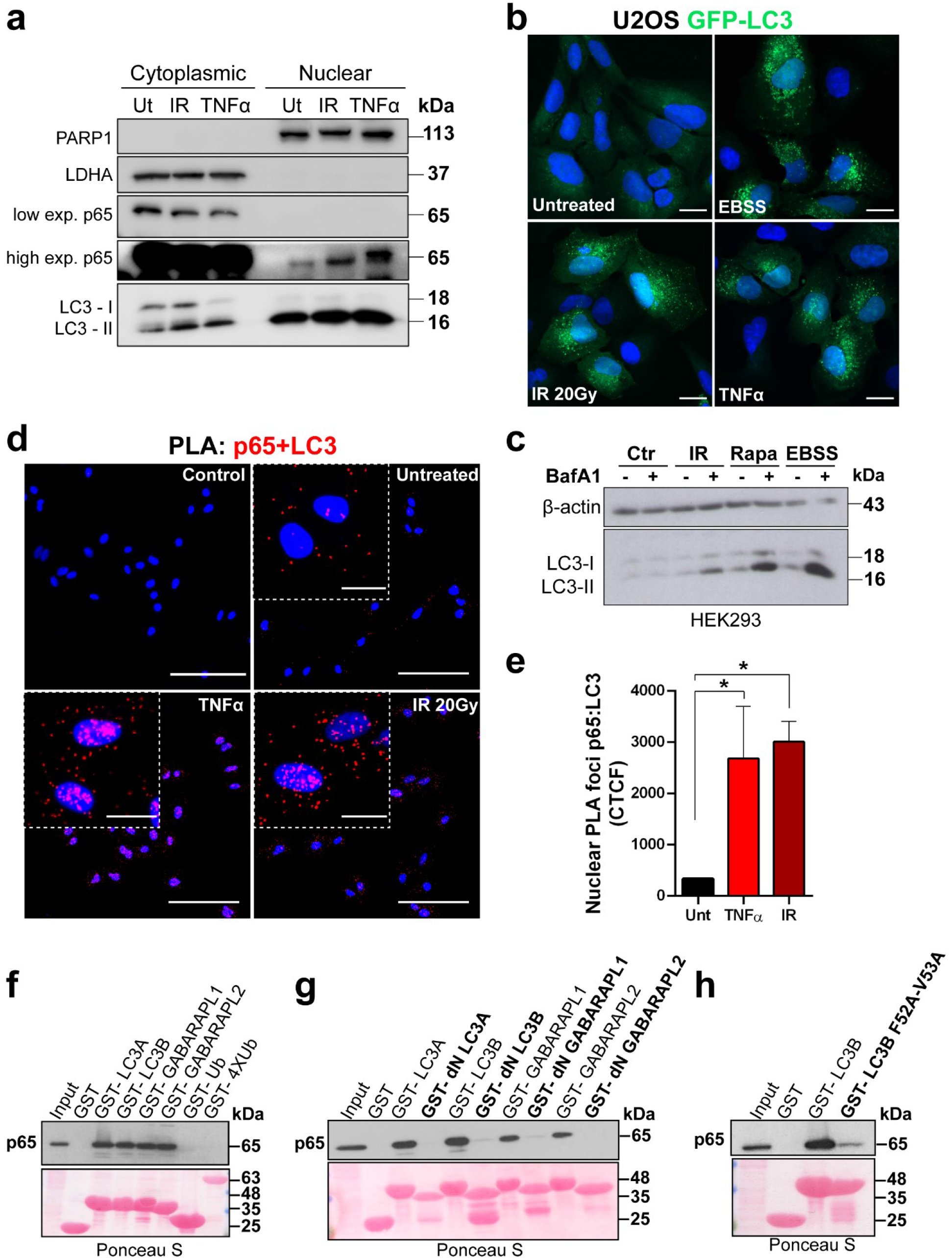
Nuclear LC3 and p65 interaction following stress-induced NF-κB activation. **a**) Western blot (WB) analysis of cytoplasmic and nuclear fractions for the autophagy marker LC3 and NF-κB subunit p65 (low exposure, high exposure) in untreated cells (Ut), after 90 min irradiation (IR 20 Gy) or 1 hour TNFα (10 ng) treatment in monocytic leukemia cells (THP1). PARP1 and LDHA are used as loading controls for nuclear and cytoplasmic fractions respectively. LC3-I, LC3-II, non-lipidated and lipidated forms of LC3. Data are representative of n=3 independent experiment. **b**) U2-OS osteosarcoma cells stably expressing GFP-LC3 were used in a GFP-LC3 puncta formation assay and analyzed in untreated cells, 1 hour after irradiation (IR 20 Gy) or TNFα treatment. Earle’s Balanced Salt Solution (EBSS) was used as autophagy positive control for starvation treatment. Images are representative fluorescence confocal microscopic photographs of n=3 independent experiments (scale bars: 20 µm). **c**) HEK293T cells were pre-treated with Bafilomycin A1 (200 nM) or DMSO (vehicle) for 2 hours and analyzed before (Ctr: control) or 1 hour after irradiation (IR 20 Gy) by WB. Rapamycin (Rapa) and starvation (EBSS) treatments were used as positive controls for autophagy induction. All data are representative of n=3 independent experiments. **d**) Proximity Ligation Assay (PLA) was used to demonstrate binding between LC3 and p65 in untreated conditions or after TNFα (10 ng) stimulation or irradiation (IR 20 Gy) treatments using A549 adenocarcinoma cells. Cells stained only with anti-p65 primary antibodies were used as a technical negative control (Control). Each red spot represents a single interaction. Nuclei were stained with DAPI. Scale bars: 100 µm and 20 µm (insets). **e**) Quantification of corrected total cell fluorescence (CTCF) of nuclear p65:LC3 PLA foci from (d). Data are means ± SEM of three independent experiments. Statistical analyses were performed by one-way ANOVA followed by Bonferroni’s multiple comparison test (*p < 0.05) using GraphPad Prism 8. **f-h**) HEK293T cell lysates were added to beads with immobilized GST fusion LC3-like modifiers (GST, GST-LC3A, GST-LC3B, GST-LC3C, GST-GABARAP, GST-GABARAPL1, GST-GABARAPL2, GST-Ub, GST-4XUb (f) or the LC3 like modifiers lacking the unique N-terminus (dN) (g) and GST-LC3BF52A-V53A (h)), followed by WB using an antibody against endogenous p65. Ponceau stain gels (bottom) represent the amount of GST beads constructs used for the GST-pulldown assays. All data are representative of n≥3 independent experiments.

Next, we studied the interaction between LC3-related proteins and p65 by performing *in vitro* pull-down experiments using GST-tagged ATG8-family members (LC3A, LC3B, and GABARAPs) as affinity baits. Endogenous p65 and overexpressed myc-tagged p65 were captured from cell lysates by all four LC3-like modifiers, but not by mono- or tetra-ubiquitin (negative controls) indicating that p65 does not possess ubiquitin-binding abilities (Fig. 1f and S1d). To further define the interaction between p65 and LC3s/GABARAPs, we analyzed LC3 mutants. Endogenous p65 did not bind to LC3 protein family members lacking the unique amino-terminal region or to LC3B mutated in the LIR docking site (LDS) which is important for recognition and interaction mechanisms (Fig. 1g, h) ^30,31^. The same results were obtained with overexpressed myc-tagged p65 (Fig. S1e, f). Together, these data indicate that the binding between p65 and LC3 family members is mediated by LC3-interacting region (LIR) motifs that bind to the LDS of ATG8 proteins

### LC3s/GABARAPs interact with the Rel homology domain of the NF-κB/p65 subunit through the cargo receptor SQSTM1/p62

In order to characterize the binding site of p65 that interacts with ATG8 proteins, we used C-terminal and N-terminal deletion mutants of p65 and performed pull-down analysis with GST-LC3B (Fig. 2a, b). As shown in Figure 2a, LC3B interacts with the Rel homology domain (RHD) of p65 between amino acids 193 and 306, a region responsible for mediating its specific DNA binding to the NF-κB consensus sequence as well as to form heterodimers with other NF-κB-family members (Fig. 2a, b). Bioinformatics analysis of the p65 protein sequence revealed the presence of six putative LIR motifs in the region of interest for LC3B interaction (https://ilir.warwick.ac.uk, ^33^). The substitution of the two hydrophobic amino acids within the LIR motif is sufficient to ablate the interaction with ATG8-family members ^34^. However, none of the single p65 LIR mutants completely lost the interaction with LC3s/GABARAPs (Fig. S2a). Degradation of selective material (cargo) can also be mediated by specific cargo receptors that can recognize damaged organelles, pathogens, or over-abundant proteins and aggregates ^35^. Many of these receptors present a ubiquitin-associated (UBA) domain through which they bind to the ubiquitinated cargo and an LIR motif to associate with the autophagosome membrane for subsequent degradation. The absence of N-ethylmaleimide (NEM), an inhibitor of deubiquitinases (DUBs) from the protein lysates made it possible to evaluate whether the binding between p65 and LC3 is ubiquitin-mediated. While NEM was included in the IP–buffer by default (Fig. 1f-h, S1d-f and Fig. 2a), when testing its requirement for interaction, we found that GST-LC3B can bind to myc-p65 only in the presence of NEM (Fig. 2c). This strongly implicates that their interaction depends on ubiquitin-conjugation.

**Fig. 2:**
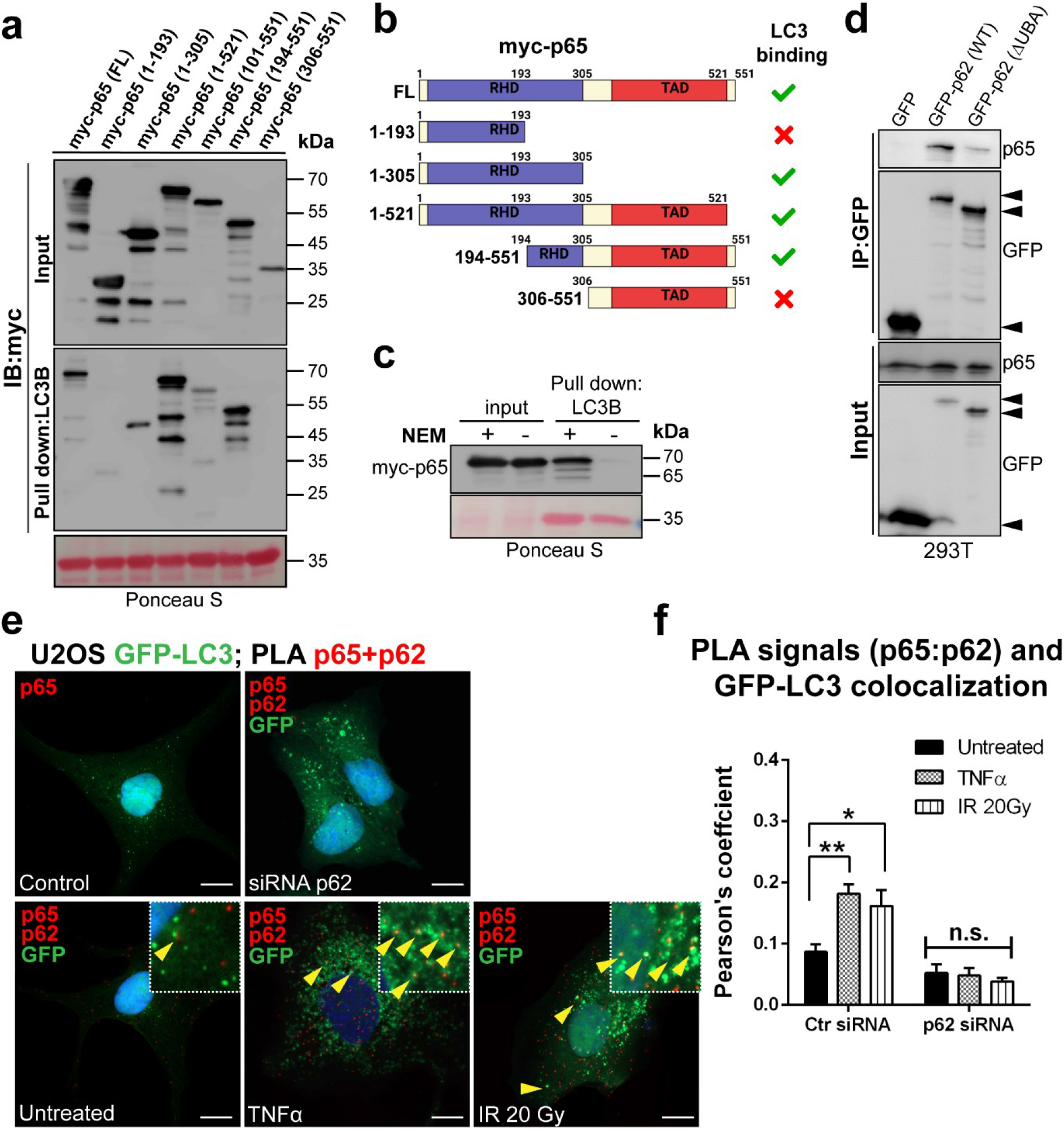
LC3 binds to the RHD of the NF-κB/p65 subunit through the ubiquitin cargo receptor p62. **a**) Full-length myc-p65 and truncation mutants have been transfected into HEK293T cells. Cell extracts were added to beads with immobilized GST-LC3B and immunoblotted for myc. LC3B was able to bind only to full-length myc-p65 and to the truncation mutants including residues 193 to 306. Ponceau stain gels (bottom) represent the amount of GST beads constructs used for the GST-pulldown assays. All data are representative of n≥3 independent experiments. **b**) Schematic representation of myc-p65 full length (FL) and truncation mutants indicating which bind to LC3. **c**) HEK293T cell lysates were added to GST-LC3B in presence or absence of NEM (N-ethylmaleimide). Overexpressed myc-p65 binds to LC3B only in the presence of NEM. All data are representative of n=3 independent experiments. **d**) HEK293T cells were transfected with empty GFP vector (control), WT, or ΔUBA deleted mutant of GFP-tagged p62. After 24 hours, cell lysates were immunoprecipitated with GFP-Trap beads and immunoblotted for GFP fusion proteins and p65. (**e**) PLA between p65 and p62 (red spots) was performed in U2-OS cells stably expressing GFP-LC3 (green spots). Top panels are representative images of cells treated only with p65 antibody (top left panel), or cells after transfection with a mixture of two siRNAs targeting p62 (top right panel; see Fig. S3b for p62 siRNA knockdown efficiency), which were used as negative controls. Bottom panels are representative images of PLA performed in untreated, TNFα (10 ng) stimulation or irradiation (20 Gy) conditions. Each red spot represents a single interaction of p62 and p65 proteins. Yellow arrowheads point to p65-p62 interacting proteins (PLA red signals) that co-localize with LC3 puncta structures (autophagosome in green). Nuclei were stained with DAPI. Scale bars 10 µm. (**F**) Pearson’s Correlation Coefficients for the colocalization of PLA signals (p65:p62) with GFP-LC3 from (e) quantified using JACoP plugin of ImageJ ^40^. Data are means ± SEM of three independent experiments (n≥5 cells for each condition). Statistical analyses were performed by two-way ANOVA followed by Bonferroni’s multiple comparison test (*p<0.05; **p < 0.01; n.s.: not significant) using GraphPad Prism 8.

The p62 protein, also called sequestosome 1 (SQSTM1), is one of the major cargo receptors able to deliver ubiquitinated proteins to the autophagosome for degradation ^36^. Moreover, previous studies proposed that p62 regulates signal transduction in NF-κB pathways as a protein adapter through the interaction with TRAF6 and RIP1 ^37,38^. However, its potential role in binding and targeting polyubiquitinated NF-κB subunits to the autophagosome during stress conditions needs to be addressed. Therefore, to examine whether LC3 binds to p65 through the cargo receptor p62, we performed immunoprecipitations after transient overexpression of p62-GFP wild-type (WT) or of a p62 mutant that lacks the UBA domain involved in binding to ubiquitinated substrates (ΔUBA) ^39^.Endogenous p65 was indeed able to bind to full-length p62-GFP (WT), but only in a greatly reduced manner to the ΔUBA mutant, confirming that their interaction is promoted by ubiquitin conjugation (Fig. 2d). In addition, PLA analysis using U2-OS cells stably expressing GFP-tagged LC3 showed a significant increase of p65-p62 binding at the level of the autophagosome membrane in the proximity of the nucleus under stress conditions (see yellow arrowheads; Fig. 2e, f and S2d, S3b). Accordingly, immunoprecipitation and immunofluorescence analysis revealed stronger p65-p62 interaction localized in the cytoplasm, in the nucleus and at peri-nuclear positions following TNFα stimulation or irradiation (Fig. S2b, S2c). Collectively, these findings suggest that selective autophagy can control the nuclear level of p65 after NF-κB activation through p62 binding.

### Inhibiting autophagy influences early stress-induced NF-κB-dependent gene expression and impairs p65 degradation

Since our findings indicated that the interaction of p65 with p62 was enhanced after induction of NF-κB by TNFα or irradiation (see 2e and S2B), we asked whether autophagy can influence the transcriptional activity of NF-κB-dependent genes. To do so, we used a transcription activity-based luciferase assay to monitor the NF-κB signal transduction pathway (NF-κB/293/GFP-Luc™, System Biosciences). The NF-κB reporter (luc)-HEK293 cell line was treated with TNFα or irradiation in the presence (or absence) of chloroquine, an inhibitor of autophagy flux by decreasing autophagosome-lysosome fusion and degradation ^24^ (Fig. 3a). Cells were transfected with renilla luciferase reporter vector 24 hours before treatment for data normalization. Results showed a significant increase in NF-κB luciferase activity after TNFα or irradiation treatment in presence of chloroquine compared to its absence (Fig. 3a, left panel), suggesting that autophagy degradation impairment can modulate the NF-κB response. To assess this hypothesis, we treated the NF-κB reporter cell line using the same stress conditions (TNFα or irradiation), but this time in the presence (or not) of the mTORC1-selective inhibitors rapamycin, which is consequently able to hyperactivate autophagy ^24^ (Fig. 3a). As expected, rapamycin significantly reduced NF-κB luciferase activity (Fig. 3a, left panel).

**Fig. 3:**
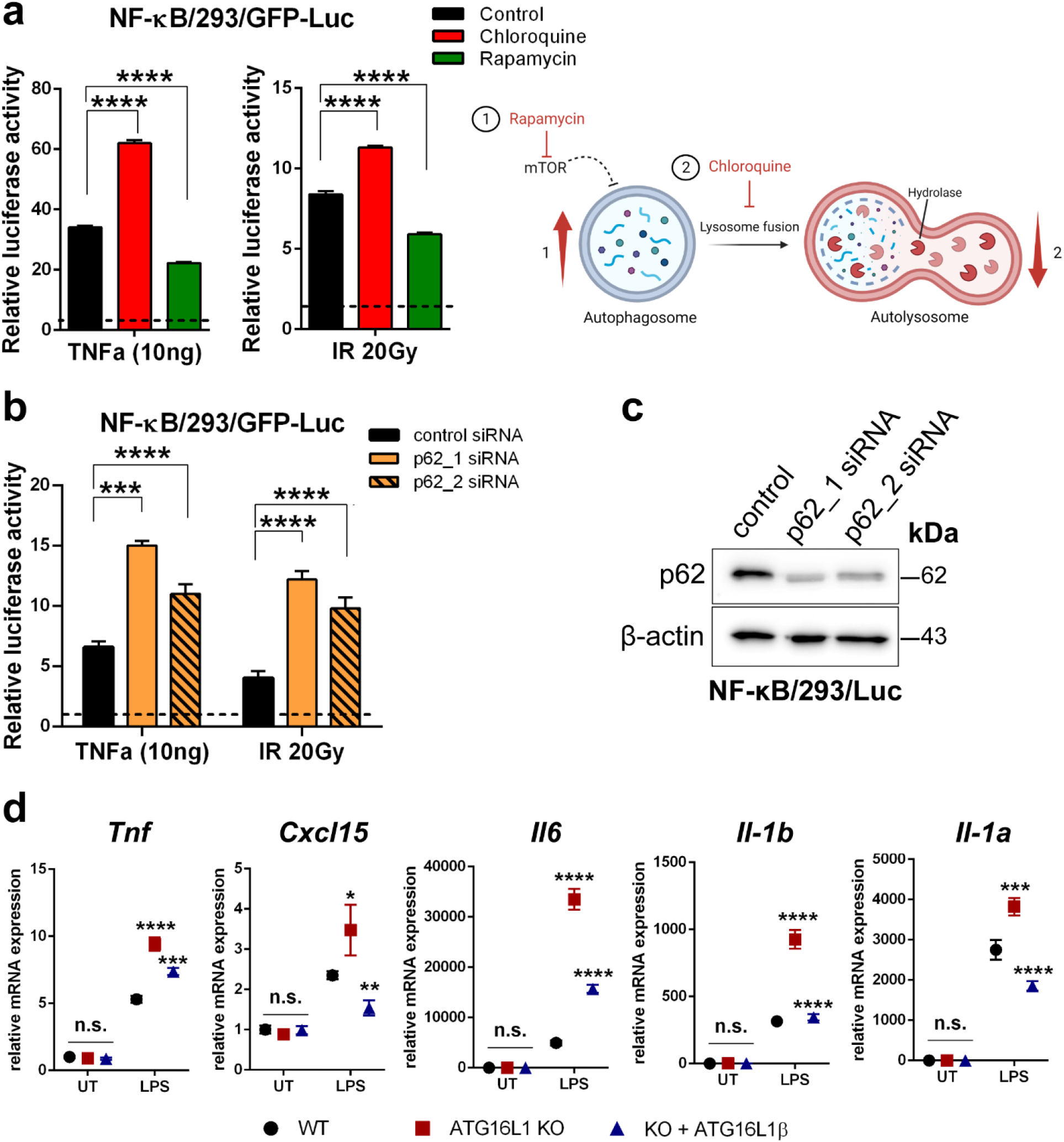
Autophagy degradation restricts the inflammatory response of NF-κB. **a, left**) The NF-κB reporter (luc)-HEK293 cell line was treated with TNFα (10 ng) or irradiation (20 Gy) in the presence or absence of chloroquine (60 µM) or rapamycin (1 μM). Cells were harvested 5 h after treatment and were subjected to luciferase activity analysis. Renilla luciferase reporter vector was used as internal control. Data are means ± SEM of seven independent experiments. Statistical analyses were performed by one-way ANOVA followed by Bonferroni’s multiple comparison test (****p<0.0001) using GraphPad Prism 8. **a, right**) Schematic representation of rapamycin (1) and chloroquine (2) inhibitory functions. Illustrations were created with BioRender.com. **b**) The NF-κB reporter (luc)-HEK293 cell line was treated with TNFα (10 ng) or irradiation (20 Gy) 48 hours after siRNA transfection. Cells were harvested 5 hours after treatment and were subjected to luciferase activity analysis. Renilla luciferase reporter vector was used as internal control. Results are normalized to the unstimulated controls, respectively. Data are means ± SEM of three or more independent experiments. Statistical analyses were performed by two-way ANOVA followed by Bonferroni’s multiple comparison test (***p<0.001, ****p<0.0001) using GraphPad Prism 8. **c**) Western blot analysis of p62 confirming the siRNA knockdown efficiency. β-actin is used as loading control. **d**) mRNA expression of *Tnf, Cxcl15* (also known as *Il8*), *Il6, Il-1β* and *Il-1α* was detected by real-time RT-qPCR in *WT* and *ATG16L1 KO* RAW 264.7 cells rescued or not with *ATG16L1β* (*KO* + *ATG16L1β*) and treated with LPS (10 µg). Results are normalized to untreated *WT* cells (UT) as controls. Data are means ± SEM of three independent experiments. Statistical analyses were performed by two-way ANOVA followed by Bonferroni’s multiple comparison test (statistical significance showed between *WT* vs *ATG16L1 KO* and *ATG16L1 KO* vs *KO* + *ATG16L1β;* *p < 0.05, **p<0.01, ***p<0.001, ****p<0.0001) using GraphPad Prism 8.

It has been suggested that activation of NF-κB through persistent impairment of autophagic degradation is a consequence of p62 accumulation ^41^. However, under conditions where stress-induced-autophagic degradation was active (see Fig. 1c), TNFα or irradiation did not induce *p62* mRNA expression (Fig. S3a). Thus, the choice to study NF-κB activation at early time points, *i*.*e*. a few hours, excludes any effect that the accumulation of p62 protein at more extended times of autophagy inhibition could exert ^41^. In addition, using different inducers, like TNFα or genotoxic stress, which stimulate different NF-κB activation pathways, would help to exclude additional factors that can influence the experimental read-out. To further explore whether autophagy influences NF-κB activity through selective binding of p62 to p65, we performed NF-κB luciferase reporter assays after silencing p62. The NF-κB reporter (luc)-HEK293 cell line was transfected with two different siRNAs against p62 before treatment with TNFα or irradiation and luciferase activity was measured as previously described. A significant increase in NF-κB luciferase activity after TNFα or irradiation treatment was observed when p62 was downregulated by both siRNAs (Fig. 3b, c), which was also reflected by the increased expression of pro-inflammatory NF-κB target genes (Fig. S3b, c).

The ATG16L1 protein is an essential autophagy component required for LC3 lipidation and autophagosome maturation ^42^. Because NF-κB plays a central role in coordinating inflammatory responses in macrophages ^43,44^, we further validated whether autophagy impacts NF-κB transcription activity using RAW264.7 cells deficient for both, α and β isoforms of ATG16L1 (*knock-out, KO*) ^45^ (Fig. 3d). LPS is a potent pro-inflammatory pathogen-associated molecular pattern that can induce both NF-κB and autophagy signaling pathways via Toll-like receptor 4 (TLR4) in macrophages ^46–48^. LPS-induced expression of NF-κB-driven inflammatory genes (*Tnf, Cxcl15* also known as *Il-8, Il6, Il-1α*, and *β*) was significantly increased in absence of *ATG16L1* compared to wild-type cells (Fig. 3d). Interestingly, stable expression only of the *ATG16L1β* isoform (*KO* + *ATG16L1β*), which is known to be sufficient to rescue the autophagic flux ^45^ in *ATG16L1 KO* cells, was sufficient to significantly reduce cytokine expression to the level comparable to *WT* macrophages (Fig. 3d). Together, our results confirm that autophagy limits stress-induced pro-inflammatory NF-κB-dependent gene expression.

Previous studies have suggested that autophagy could trigger p65 degradation by lysosomes, although the exact mechanism is still unclear ^49–51^. Confocal microscopic analysis revealed that TNFα stimulation caused enhanced p65 colocalization with the lysosome-associated membrane glycoprotein (LAMP1), a well-known lysosomal marker (Fig. 4a) ^52^. To further assess the autophagy/lysosome-dependent clearance of p65, we used bafilomycin A1, a lysosomal inhibitor. Pretreatment with bafilomycin A1 enhanced colocalization of p65 and LAMP1, as well as p65 and LC3 after TNFα stimulation (Fig. 4a). This indicates that p65 is delivered from autophagosomes to lysosomes and that this process is enhanced after activation of NF-κB. Next, we determined the turnover rates of p65 protein after TNFα stimulation by performing cycloheximide (CHX)-chase experiments (Fig. 4b, c). To avoid the influence of p65 expressed asynchronously on the measurements of protein turnover, cells were transfected 24 hours before treatment with myc-p65. Subsequently, cells were stimulated for 15 min with TNFα and chased for four-time points (0-6 h) by cycloheximide addition in a fresh medium. Western blot analysis showed a significant myc-p65 reduction at 6 hours and a half-life between 3 and 6 hours (Fig. 4b, c). Moreover, immunoprecipitation of overexpressed GFP-LC3 showed initial binding of endogenous p65 at 1 hour which increased at 3 hours and is lost at 6 hours after TNFα treatment (Fig. 4d). Saccani et al. (2004) proposed that p65 sequestered to its specific DNA binding sites is degraded by the proteasomal pathway ^10^. In order to clarify if autophagy also plays a role in regulating the protein stability of p65 after TNFα stimulation, we performed a degradation assay using a proteasomal (MG132), or an autophagy-lysosome (chloroquine, CQ) inhibitor (Fig. 4e). Surprisingly, myc-p65 protein degradation further declined when MG132 was present after TNFα washout either at 6 or 24 h (Fig. 4e). In contrast, chloroquine was able to rescue its protein level over time (Fig. 4e). It is worth noting that exposure of cells to MG132 caused markedly increased autophagy as shown by the conversion of LC3-I to lipidated LC3-II at 6 h and sustained up to 24 h (Fig. 4e), which could explain the further decrease of p65 protein level compared to control after TNFα washout. These data confirmed autophagy as an alternative mechanism responsible for p65 degradation to control NF-κB activation.

**Fig. 4:**
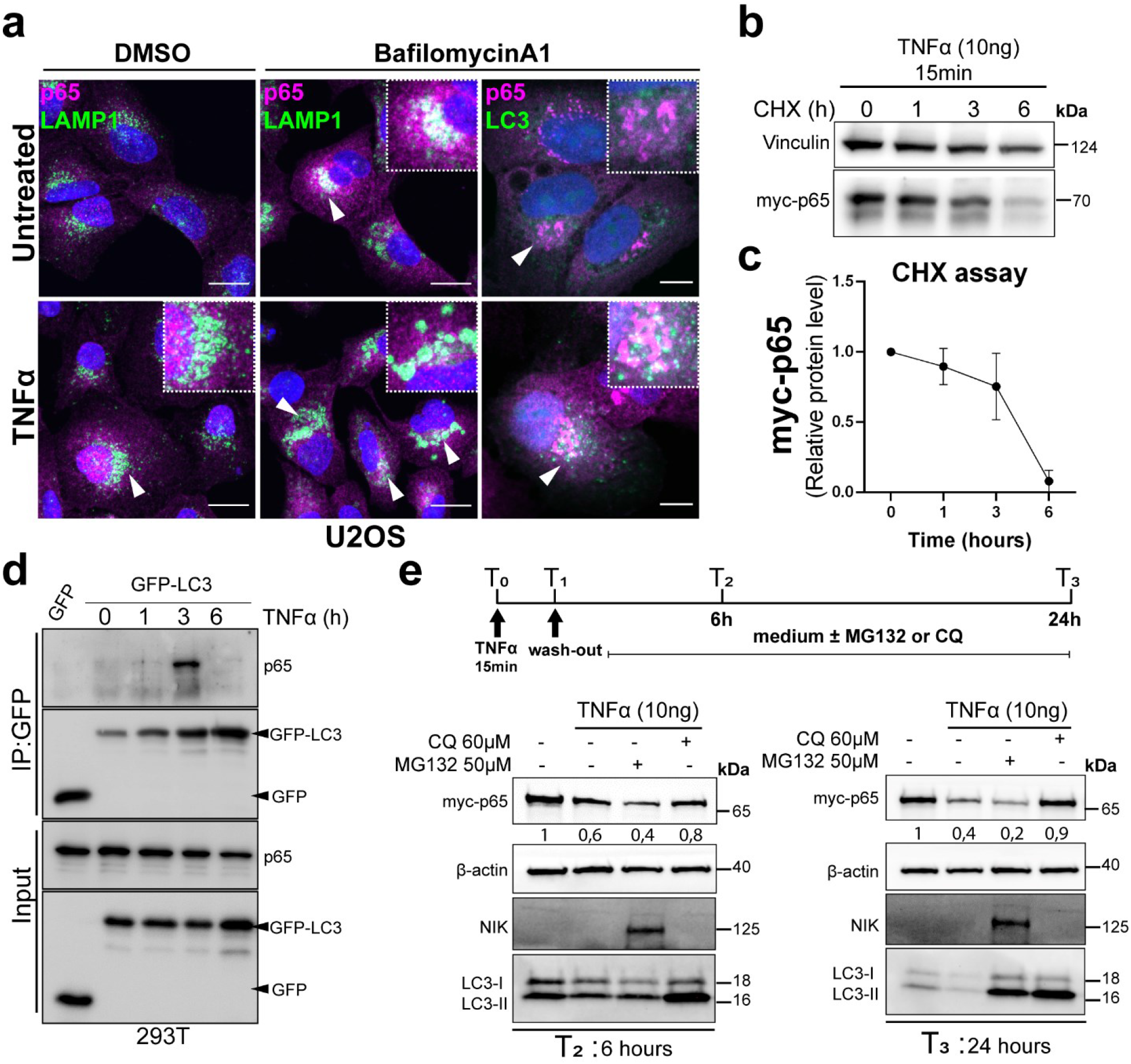
Lysosomal degradation of p65 after NF-κB activation. **a**) Immunofluorescence staining for p65 (magenta) and LAMP1 or LC3 (green) using U2-OS cells pre-treated 30 minutes with bafilomycin A1 (200 nM) or DMSO (control) in untreated or TNFα (10 ng) stimulated conditions. White arrowheads indicate colocalization of p65 in the lysosomes (LAMP1) or autophagosome (LC3) compartments. DAPI was used for the staining of nuclei. Z-stack images shown are representative fluorescence confocal microscopic photographs of n=3 independent experiments (scale bars 20 µm). **b and c**) Cycloheximide-chase assay performed in HCT116 transfected with myc-p65 24 hours before treatment. Cells were washed once after 15 minutes of TNFα stimulation and cycloheximide was added. The time course was performed for a total of 6 hours as shown in the graph and analyzed by western blot (**b**) at the times indicated using an antibody against myc. Vinculin is used as loading control and quantification of the protein abundance was done using ImageJ (**c**). Data are representative of n=3 independent experiments. **d**) HEK293T cells were transfected with GFP empty vector (control) or GFP-LC3. After 24 hours, cells were treated with TNFα (10 ng) and cell lysates were collected at the time points indicated. Immunoprecipitation was performed with GFP-Trap beads and samples immunoblotted for GFP fusion protein and p65. (**e**) HCT116 were transfected with myc-p65 plasmid 24 hours before the 15 minutes TNFα treatment. Cells were washed once with PBS and fresh medium was added for 6 hours (T_2_, left blot) or 24 hours (T_3_, right blot) with or without MG132 (10 µM) or chloroquine (CQ, 60 µM). On top is a schematic representation of the experimental setup used. Western blot analysis was performed using antibodies against myc and β-actin as loading control. Antibodies against NIK and LC3-I/II were used as controls for MG132 and CQ inhibitors respectively. All data are representative of n=3 independent experiments.

### Autophagy affects the strength and duration of NF-κB activation by controlling p65 nuclear occupancy

In addition to the robust mechanism of inhibition mediated by IκBα, NF-κB-induced transcription is selectively controlled in a gene-specific manner by post-translational modifications ^10^. In this regard, ubiquitination of p65 occurs predominantly in the nucleus which controls DNA binding of a fraction of nuclear p65 ensuring the limited duration of expression of specific NF-κB target genes ^10,53^. A study based on transcriptome analysis and NF-κB dynamics concluded that increased expression of inflammatory cytokine genes in macrophages requires sustained p65 nuclear occupancy ^54^. We showed that accumulation of LC3 and p62 within the nucleus strongly co-localized with p65 following NF-κB activation (see Fig. 1d, S1c, and S2b). Moreover, the mRNA expression of various pro-inflammatory NF-κB target genes was significantly higher in macrophages upon LPS stimulation when autophagy degradation was impaired (Fig. 3d). Thus, we asked if autophagy impacts the dynamics of NF-κB/p65 after activation. To this end, the nuclear translocation of p65 after LPS treatment was measured in a time-resolved manner. Western blot analysis indicated significantly increased and prolonged permanence of p65 in the nucleus at 3 and 6 hours after LPS stimulus in autophagy knock-out macrophages (*16L1 KO*) compared to the wild-type control (*WT*) (Fig. 5a, b). Accordingly, hyper-activation of NF-κB was also shown at 3 hours after LPS in *ATG16L1 KO* macrophages compared to control, confirming our findings (Fig. 5c).

**Fig. 5:**
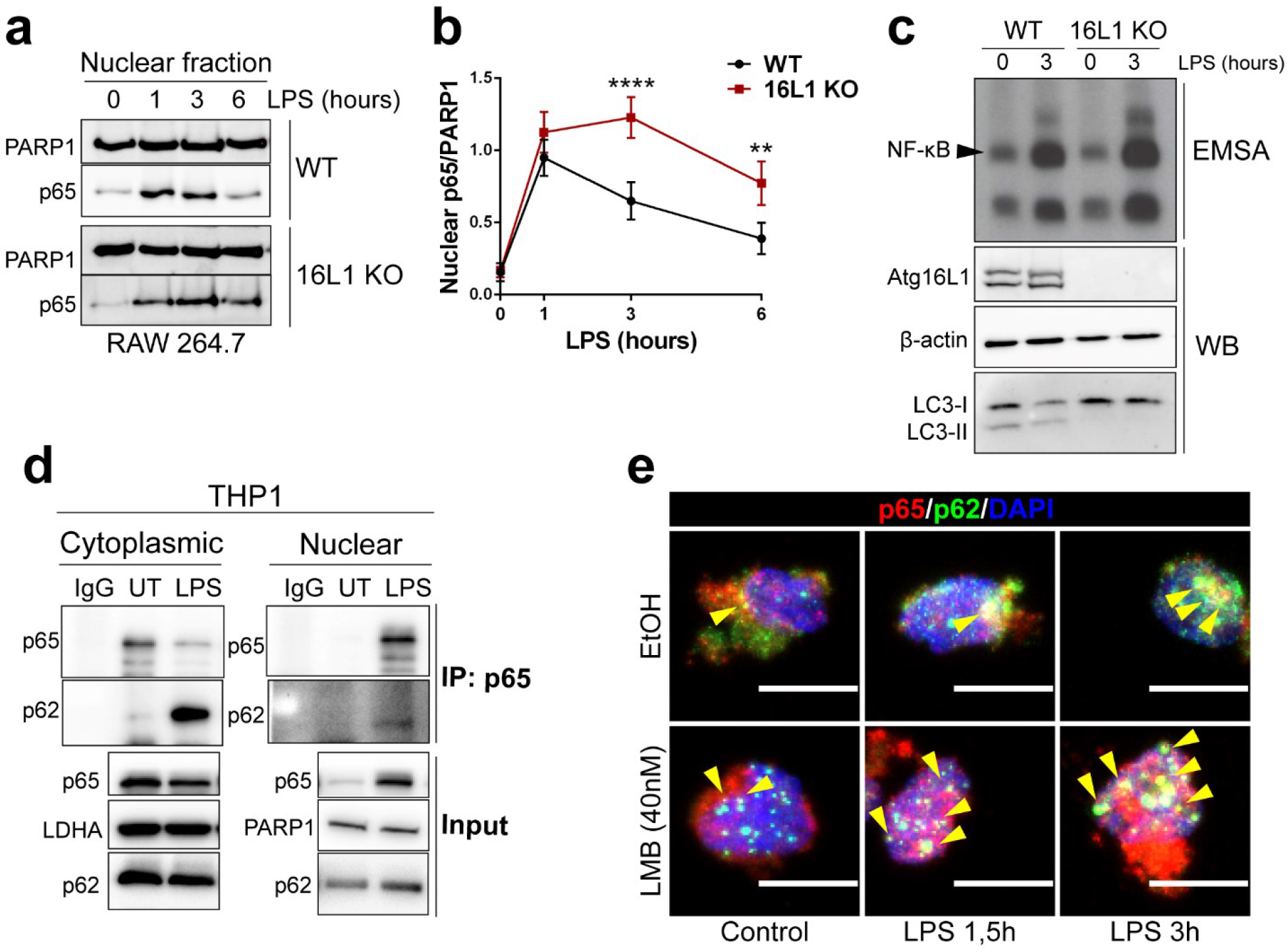
Autophagy impairment affects p65 nuclear retention. **a, b**) Nuclear extracts from *WT* and *ATG16L1* KO RAW 264.7 cells were collected at the time points indicated after LPS (10µg) treatment and analyzed by WB (a) for p65. PARP1 is used as nuclear loading control. Quantifications of the protein abundance (b) were measured using ImageJ and statistical analysis was performed by Two-way ANOVA followed by Bonferroni’s multiple comparison test (**p<0.01, ****p<0.0001) using GraphPad Prism 8 (n=3). **c**) *WT* and *ATG16L1* KO RAW 264.7 cells were treated with LPS for the times indicated and whole cell extracts were analyzed by EMSA for NF-κB activity (top panel) or immunoblotted (WB) with the indicated antibodies (bottom panel). Data are representative of n=3 independent experiments. **d**) Cytoplasmic and nuclear fractions from THP1 cells were extracted at 0 or 90 min after LPS (10 µg) treatment for immunoprecipitation (IP) with anti-p65 antibody, followed by WB analysis with anti-p65 or anti-p62 antibodies. LDHA and PARP1 antibodies were used as loading controls for cytoplasmic and nuclear fractions, respectively. IgG was used for IP control. **e**) Immunofluorescence staining for p65 (red) and p62 (green) using THP1 cells pre-treated with leptomycin B (LMB 40 nM) for 2-hours followed by LPS (10 µg) treatment for the time indicated. Yellow arrowheads indicate colocalization between p65 and p62 in the nuclei stained with DAPI. Z-stack images shown are representative fluorescence confocal microscopic photographs of n=3 independent experiments (scale bars 10 µm).

p62 has two nuclear localization signals and one nuclear export signal enabling nucleocytoplasmic shuttling, however, little is known about its possible nuclear roles ^55^. To determine if p62 controls a fraction of nuclear p65 in response to LPS treatment, immunoprecipitation of endogenous p65 was performed in cytoplasmic and nuclear fractions from THP1 cells. Although p62 co-immunoprecipitated with p65 mainly in the cytoplasm, a small quantity of their binding was also found in the nucleus after LPS-induced NF-κB activation (Fig. 5d). Nucleocytoplasmic shuttling of p62 before (Control) and after LPS treatment was further shown using Leptomycin B (LMB), which is a potent and specific nuclear export inhibitor (Fig. 5e) ^56^. Immunofluorescence staining of p65 and p62 in THP1 cells revealed that their nuclear colocalization upon LPS treatment increased when the nuclear export was inhibited by LMB (Fig. 5e), suggesting that p62 could bind to p65 in the nucleus and translocate it to the cytoplasm together with LC3 where they are subject to autophagic degradation. Collectively, our data demonstrate a novel role for the cargo receptor p62 in trafficking ubiquitinated p65 from the nucleus to autophagosomes for degradation to control inflammation-driven NF-κB hyperactivation.

## Discussion

Autophagy plays a critical role in inflammation and immunity, as evidenced by previously identified correlations between autophagy-related gene polymorphisms and many inflammatory disorders, including cancer ^18,19,57,58^. For instance, several autophagy-deficient mice models, such as myeloid-specific *Atg16L1* KO mice, colonic epithelial cell-specific *Atg7* KO mice, and *Atg4B* null mice all exhibit exacerbated colitis induced by dextran sodium sulfate (DSS) ^59–61^. Moreover, autophagy intrinsically regulates the function of the immune system. Inhibition of macrophage autophagy promotes inflammatory M1 macrophage polarization which is characterized by excessive cytokine production - a phenotype also observed in aged macrophages with a significantly decreased autophagy level ^49,61,62^. All these studies suggest that impaired autophagy promotes a pro-inflammatory condition. However, despite the increasing interest in the role of autophagy in the pathogenesis of several autoimmune and inflammatory disorders, the mechanism by which autophagy prevents exaggerated pro-inflammatory responses needs further investigation. Hence, understanding the exact molecular mechanism that facilitates pro-inflammatory responses upon inhibition of autophagic activity greatly impacts the development of new therapeutic strategies for the treatment of various disorders.

The NF-κB signaling pathway and autophagy are both implicated in determining cellular fate, influencing each other through positive or negative feedback loops ^63^. Recent evidence already suggested that autophagy suppresses NF-κB signaling to limit the inflammatory response. Constitutive activation of NF-κB/p65, leading to increased expression of pro-inflammatory cytokines, was found associated with autophagy suppression either after kidney injury or in intestinal epithelial cell (IEC)-specific *Atg7* KO mice ^20,23^. By using biochemical analysis and imaging techniques we characterized the interaction of endogenous autophagic proteins and the NF-κB/p65 subunit upon stress-induced inflammation. Strikingly, accumulation of LC3 within the nucleus strongly co-localized with p65 following NF-κB activation, indicating that nuclear LC3 sequesters p65 protein from the nucleus to the cytoplasm to block NF-κB signaling. Previous studies showed that autophagy may degrade nuclear components, but the molecular targets and their underlying mechanism in the nucleus remained incompletely understood ^25^.

Nuclear accumulation of LC3 plays a role in different nuclear functions in response to specific types of stress ^27,64,65^. In addition to LC3, autophagosome proteins ATG5 and ATG7 have also been identified in the nucleus regulating p53 activation ^66,67^. Similar to p53, ATG5 was found to co-localize with p65 in the nucleus in renal epithelial cells to alleviate tubular cell inflammation in response to kidney injury induced by unilateral ureteric obstruction (UUO) or angiotensin II (Ang II) ^20^. However, the molecular mechanism involved in ATG5-p65 complex formation was not fully elucidated. Our study is the first to show that the activity of a transcription factor can be directly regulated by selective autophagy in the nucleus to prevent exaggerated pro-inflammatory responses. We demonstrate that the p65-LC3 interaction is promoted by ubiquitination of the same p65 protein, which is recognized by p62.

p62 is a well-known autophagy cargo receptor that recognizes the ubiquitinated proteins for selective degradation via its ubiquitin-binding domain (UBA) and delivers them to autophagosomes via its LC3 interaction region (LIR) ^36^. Interestingly, Lobb et al. (2021) recently described a novel function for p62 in trafficking nuclear-ubiquitinated p65 to nucleolar aggresomes in response to aspirin, which induced apoptosis in cancer cells ^68^. Furthermore, they suggested an active competition between the autophagic and nucleolar accumulation of the protein, which might partially explain the difference found in our study. Another explanation may be the different stimuli used for NF-κB induction. For instance, the nucleolar trafficking of p65 is restricted to specific stimuli such as aspirin and proteasome inhibitors, while nucleolar sequestration of p65 in response to TNFα, LPS, or γ-irradiation has not been previously reported ^69^. Although the exact signals that regulate the nuclear-cytoplasmic shuttling of p65-p62 remain to be evaluated in the future, the present study is a direct proof that p62 recognizes nuclear-ubiquitinated p65 and limits the duration of NF-κB activity.

Adequate control of the termination of the NF-κB response is of fundamental importance to prevent sustained production of inflammatory mediators and thus immunopathology such as inflammatory bowel disease (IBD) and cancer ^70,71^. Direct inhibition of NF-κB family member p65 abrogates established experimental colitis, underlining the importance of p65 in chronic intestinal inflammation and the need for an effective therapeutic strategy able to repress its activity ^72^. In contrast to the increased knowledge of the mechanisms leading to NF-κB activation, little is known about the inhibitory mechanisms regulating its inactivation. Here, we demonstrated that p65 ubiquitination and autophagosomal degradation is a mechanism of post-transcriptional repression to control inflammation-driven NF-κB hyperactivation. Inhibition of autophagy activity by depletion of an essential autophagy gene *ATG16L1* selectively stabilizes nuclear p65, which in turn enhanced the expression of several pro-inflammatory cytokines. The gene *ATG16L1* is one of the most important susceptibility genes associated with the pathogenesis and progression of IBD ^58^. The IκBα encoding gene *NFKBIA*, was also identified as a risk locus for IBD, suggesting that both autophagy defects and reduced levels of IκBα might contribute to hyperactive nuclear NF-κB which correlates with the severity of intestinal inflammation ^73,74^. Accordingly, Matsuzawa-Ishimoto et al. (2017) recently demonstrated that ATG16L1 in the intestinal epithelium is essential for preventing loss of Paneth cells and reducing TNFα-induced IBD pathology both *in vivo* and *ex-vivo* ^75^. Similarly, depletion of Paneth cells was also observed in the small intestine of mice with constitutive activation of NF-κB, which induces several hallmarks of IBD including increased apoptosis and mucosal inflammation ^70^. Moreover, increased proteasomal degradation determines higher rate proteolysis of IκBα protein in the mucosa of IBD patients ^76^. We also observed that autophagy affects NF-κB-driven inflammatory responses in epithelial cells upon irradiation induced-genotoxic stress. Patients with IBD are at an increased risk of developing colorectal cancer later in life ^1^. Therefore, our data imply that in the subset of IBD patients with missense polymorphisms in the *ATG16L1* gene and thus reduced autophagy, an alternative to the traditional radiation therapy should be considered.

In conclusion, we reveal a novel molecular mechanism modulating the NF-κB inflammatory response through nuclear selective sequestration of p65 by autophagy proteins. Therefore, the re-establishment of autophagy flux might alleviate the inflammatory response through selective inhibition of NF-κB signaling. Given the critical role of NF-κB in chronic inflammatory diseases (for example Crohn’s Disease and Ulcerative Colitis) and cancer (Colorectal Carcinoma), our findings are of great importance for future translational studies focusing on the modulation of its activity with great clinical impact for the prevention and/or treatment of inflammatory disease.

## Experimental Procedures

### Cell lines and reagents

All cell lines were grown under sterile and standard cell culture conditions (5% CO_2_ at 37 °C) and routinely tested for mycoplasma. The NF-κB/293/GFP-Luc™, 293T, U2-OS, A549, RAW 264.7, HCT 116, and MEF cells were grown in high glucose-containing Dulbecco’s Modified Eagle Medium (DMEM, Gibco) and were supplemented with 10 % FBS (Gibco) and 1% penicillin/streptomycin (Gibco). THP1 cell lines were grown in Roswell Park Memorial Institute Medium 1640 (RPMI, Gibco) supplemented as described above. The *ATG16L1* knockout, wild type, and rescued HEK293 and RAW 264.7 cells were kindly provided by Alf Håkon Lystad and Anne Simonsen ^45^. The U2-OS cells stably expressing GFP-LC3 were a kind gift from Ivan Dikic.

The following reagents were used for cell treatments at indicated concentrations: 10 ng/ml TNFα (Enzo Life Science), 10 µg/ml LPS (Sigma), 10 µM MG-132 (Enzo Life science), 200 nM of Bafilomycin A1 (Enzo Life science), 60 µM of Chloroquine (Enzo Life science), 1 µM of Rapamycin (Selleck), 40nM Leptomycin B (Cell signaling. Ionizing irradiation (20 or 80 Gy, as indicated) of cells was applied with a Cs137 source (OB29 Irradiator, STS Braunschweig). Cells were starved using Earle’s Balanced Salts medium (EBSS) (Sigma) for 4 hours.

### Antibodies, plasmids, and siRNA

All primary and secondary antibodies for immunoblotting were used at 1:1,000 and 1:5,000 dilutions, respectively. The following antibodies were used for the experiments in this study: anti-IκBα (clone C-21; sc-371), anti-LDHA (clone N-14; sc-27230), anti-PARP1 (clone F2; sc-8007), anti-p50 (sc-114p) were from Santa Cruz; anti-b-actin (13E5; 49970), anti-NF-κB p65 (D14E12; 8242); anti-phospho-NF-κB p65 (93H1; 3033) were from Cell Signaling; anti-LC3B (NB100-2220) was from Novus Biologicals; anti-Vinculin (V9131), anti-myc (clone 9E10; M5546) were from Sigma; anti-GFP (ab6673) and anti-LAMP1 (ab24170) were from Abcam; anti-ATG16L1 mouse mAb (M150–3) was from MBL, anti-p62 (PW9860) was from Enzolife and anti-p62 (M01; H00008878) was from Abnova. HRP-conjugated antibodies were obtained from Jackson ImmunoResearch (711-035-152, 705-035-147, and 715-035-150). Alexa-488 and −546 conjugated secondary antibodies were obtained from Invitrogen (A11008 and A21123).

Plasmid constructs are listed in Table 1. All plasmids were verified by DNA sequencing.

**Table 1.** Constructs used in this study.

| Plasmid | Description |
| --- | --- |
| pcDNA3 | Invitrogen |
| pRenilla | Addgene |
| GFP-p62-WT | p62 full-length subcloned into pEGFP-C2 <sup>39</sup> |
| GFP-p62-ΔUBA | p62 fragment subcloned into pEGFP-C2 <sup>39</sup> |
| pcDNA3-human p65 FL | myc N-terminal-tag <sup>77</sup> |
| p65 aa 1-193 | myc N-terminal-tag |
| p65 aa 1-305 | myc N-terminal-tag |
| p65 aa 1-521 | myc N-terminal-tag |
| p65 aa 101-551 | myc N-terminal-tag |
| p65 aa 194-551 | myc N-terminal-tag |
| p65 aa 306-551 | myc N-terminal-tag |
| pGEX-4T1 | GST only <sup>78</sup> |
| pGEX-4T1 LC3A dG | GST-tag <sup>78</sup> |
| pGEX-4T1 LC3B dG | GST-tag <sup>78</sup> |
| pGEX-4T1 LC3C dG | GST-tag <sup>78</sup> |
| pGEX-4T1 GABARAP dG | GST-tag <sup>78</sup> |
| pGEX-4T1 GABARAP-L1 dG | GST-tag <sup>78</sup> |
| pGEX-4T1 GABARAP-L2 dG | GST-tag <sup>78</sup> |
| pGEX-4T1 Ub | GST-tag <sup>78</sup> |
| pGEX-4T1 4XUb | GST-tag <sup>78</sup> |
| pGEX-4T1 dN LC3A dG | GST-tag <sup>79</sup> |
| pGEX-4T1 dN LC3B dG | GST-tag <sup>79</sup> |
| pGEX-4T1 dN GABARAP-L1 dG | GST-tag <sup>79</sup> |
| pGEX-4T1 dN GABARAP-L2 dG | GST-tag <sup>79</sup> |
| pGEX-4T1 LC3B F52A-V53A dG | GST-tag <sup>79</sup> |

siRNA Sequences: SQSTM1 (S_1) 5’-GGACCCAUCUGUCUUCAAA UU, SQSTM1 (AS_1) 3’-UU CCUGGGUAGACAGAAGUUU; SQSTM1 (S_2) 5’-GCCAUCCUGUUAAAUUUGU UU SQSTM1 (AS_2) 3’-UU CGGUAGGACAAUUUAAACA; ATG5_Sigma EHU085781; Non-targeting 5’-UUCUCCGAACGUGUCACGU, Non-targeting 5’-ACGTGACACGTTCGGAGAA.

cDNA mutations were generated via PCR site-directed mutagenesis according to standard protocols. Primer sequences used for LIR mutagenesis can be shared upon request.

### Nuclear and cytoplasmic fractionation

After washing with PBS, cells were incubated for 30 min on ice in three volumes Buffer A (10 mM Hepes pH 8, 1.5 mM MgCl_2_, 10 mM KCl, proteinase and phosphatase inhibitors), supplemented with 0.1% NP-40 and 1mM DTT. Samples were then centrifuged at 14,000 rpm for 30 min. The obtained supernatant containing cytoplasmic extracts was transferred to a new tube. Precipitates were washed in complete Buffer A, centrifuged for an additional 30 min at 14,000 rpm and 4°C, and resuspended in two volumes Buffer C (20 mM Hepes pH8, Glycerol 25% (v/v), 0.46 M NaCl, 1.5 mM MgCl_2_, 0.2 mM EDTA, proteinase and phosphatase inhibitors) supplemented with 0.5% NP-40. Following 30 min incubation at 4°C, lysates were centrifuged for 20 min at 14,000 rpm to remove cell debris. The supernatant nuclear extract was transferred to a new tube. Samples were stored at −80°C.

### Western blotting

For western blots, cell pellets were lysed in whole-cell extraction buffer (150 mM NaCl, 50 mM Tris pH 7.5, 1% NP-40) supplemented with proteinase and phosphatase inhibitors (Roche). The lysates were cleared by centrifugation at 14,000 rpm for 30 min at 4 °C. The Western blots were processed according to standard procedures and analyzed with a chemiluminescent imaging system (ChemiDoc Imaging Systems, Bio-Rad). Quantifications for all Western blots were carried out by densitometric analysis using Image J software.

### Immunoprecipitation

Magnetic protein G Dynabeads (Thermofisher) were washed three times with 800 µl IP Washing Buffer (150 mM NaCl, 50 mM Tris pH 7.5, 5% Glycerol). After the final wash, a 50% slurry was prepared with IP Buffer (150 mM NaCl, 50 mM Tris pH 7.5, 1% NP-40, 5% Glycerol, NEM (N-ethylmaleimide), phosphatase inhibitor and complete protease inhibitor). Cells were harvested with IP Buffer. 20 µg protein samples were prepared as input samples and 2-4 mg proteins were used for co-IP. For pre-clearing, 15 µl of the 50% slurry of Dynabeads were added to the samples and incubated for 30 min at 4°C, while rotating. DynaMag ™ magnet was used to remove the supernatant that was transferred to a new tube with an additional 15 µl of Dynabeads together with 3 µg of primary antibody and rotated overnight at 4°C. The day after, the supernatant was discarded using the magnet. Beads were washed three times with IP washing buffer to remove contamination, diluted in 2x Laemmli buffer and boiled for 5 min at 95°C. In parallel, input samples were boiled. Input and co-IP samples were analyzed by SDS-PAGE and Western blot. Co-IP using the GFP beads was performed according to the manufacturer’s protocol (ChromoTek GFP-Trap).

### GST pull-down

GST fusion LC3s, GABARAPs, and Ub proteins were kindly provided by Ivan Dikic and the expression was induced as previously described ^79^. Briefly, fusion proteins were expressed in Escherichia coli BL21 (DE3) cells in LB medium after addition of 0.5 mM IPTG and incubation at 37°C for 5 hr. Harvested cells were lysed using sonication in a lysis buffer (20 mM Tris-HCl pH 7.5, 10 mM EDTA, 5 mM EGTA, 150 mM NaCl). GST fusion proteins were immuno-precipitated using Glutathione Sepharose 4B beads (GE Healthcare). Fusion protein-bound beads were used directly in GST pull-down assays. HEK293 cells were used for pull-down of endogenous p65 or overexpressed myc-p65. Lysates were cleared by centrifugation at 14000 g for 30 minutes and incubated with GST fusion protein-loaded beads overnight at 4°C. Beads were then washed three times in lysis buffer, resuspended in 2x Laemmli buffer and boiled. Supernatants were loaded on SDS-PAGE.

### Cycloheximide-chase assay

In order to study the protein turnover of p65/NF-κB, cells were stimulated with 10 ng TNFα for 15 min, washed once with 1x PBS, and finally treated with 10 µg/ml of cycloheximide (Enzo Life science) for indicated times. Cell lysates were prepared and analyzed by western blotting.

### Quantitative real-time PCR (qRT-PCR)

RNA extraction was performed according to the manufacturer’s protocol using the RNeasy kit (Qiagen). The total RNA concentration was determined using NanoDrop. In total 1µg RNA was used for cDNA synthesis using iScript cDNA synthesis kit (Bio-Rad) according to the manufacturer’s protocol. The qRT-PCR was carried out using the CFX96 Real-Time System (Bio-Rad) and GoTaq® qPCR Master Mix (Promega). Two references were used for normalization and the fold induction of mRNA of the target of interest was calculated compared to the control sample using the ΔΔCq method. All samples were analyzed in triplicates and are shown as mean, with the error bar representing the SEM.

Primer sequences used in this study for qRT-PCR are listed in Table 2.

**Table 2.**
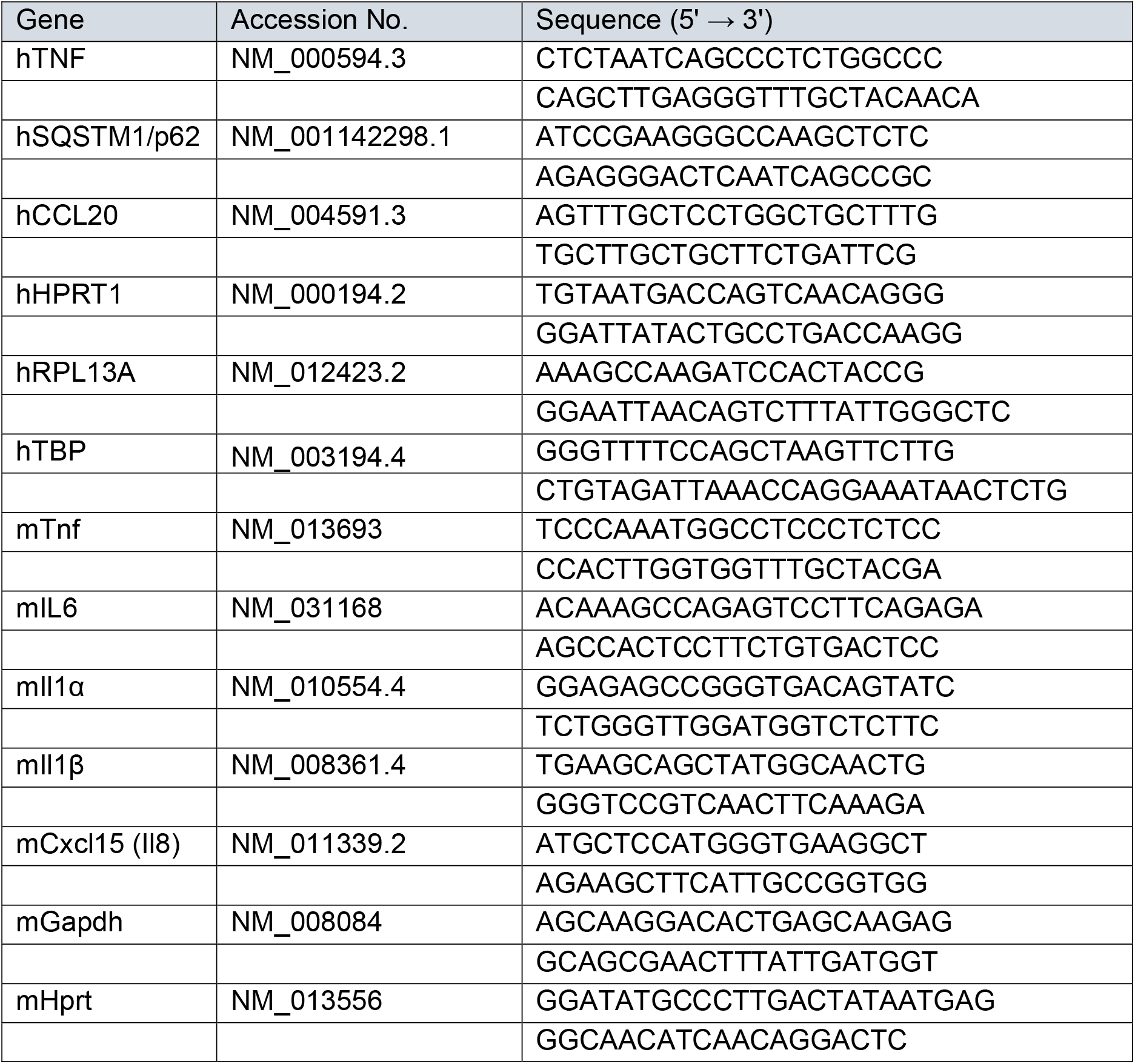
Primer sequences used in this study.

### Transfection and luciferase reporter assay

Transfection using Lipofectamine 2000 (Invitrogen) was performed according to the manufacturer’s protocol. In brief, 4 × 10^5^ cells were seeded into each well of 6-well plate 24 hours prior to transfection. Medium was replaced with DMEM without antibiotics shortly before transfection. 1 µg of DNA and 3 µl of Lipofectamine were each diluted in 150 µl OptiMEM and incubated for 5 min. The mixtures were combined and incubated for 15 min at RT. 250 µl of DNA-Lipofectamine complex was added to the cells while slowly rocking the plate followed the incubation at 37°C and 5% CO_2_. Cells were harvested 24 - 48 h following transfection.

Transient knock-down by siRNAs using Lipofectamine RNAimax (Invitrogen) was performed according to the manufacturer’s protocol. A total of 2,5 × 10^5^ cells were seeded into each well of a 6-well plate one day before the transfection. 10µM of working solutions for each siRNA were prepared. 10nM of siRNA and 9 µl of RNAimax were each added to 150 µl of OPTIMEM. The mixtures were combined and incubated at RT for 15 minutes before adding to the cells in a dropwise manner. Subsequent experiments were done 48 hours post-transfection.

The activity of the NF-κB luciferase reporter cell line was measured according to the manufacturer’s protocol (ONE-GloTM Promega) into 96-well plates with white walls and bottom to reduce the cross-contamination of the signal between wells. The Luciferase signal was measured using the plate reader (Cytation 1, Biotek).

### Electrophoretic Mobility Shift Assay (EMSA)

The EMSA was performed as previously described ^80^. In brief, 5 µg of protein extract was mixed with radioactively labeled (^32^P) H2K probes and with shift buffer (40 mM HEPES pH 7.9, 120 mM KCl, and 8% (v/v) Ficoll), 2 µg/ml poly dI/dC, 100 mM DTT and 10 µg/ml BSA and incubated for 30 min at RT. Samples were separated by electrophoresis and transferred to Whatman filter paper by vacuum dryer. The dried gel was incubated with HyperfilmTM MP at −80°C overnight and developed with standard procedures.

### Immunofluorescence analysis

Cells were grown in coverslips and fixed with 4% PFA for 10 minutes at RT, following three times PBS washing. Cells were then incubated with a quenching buffer (0.12% glycine, 0.2% saponin in PBS) for 10 minutes and incubated with a permeabilization buffer (0.2% saponin in PBS) for additional 10 minutes at RT. After three times washing with PBS, cells were incubated with a blocking buffer (10% FBS, 0.2% saponin in PBS) for 1 hour. According to the manufacturer’s dilution range recommendations, the respective antibody was diluted in the blocking solution. Coverslips were embedded in the pre-diluted primary antibody solution and incubated at 4 C° overnight. After washing with permeabilization buffer, A FITC conjugated secondary antibody recognizing the host species of the respective primary antibody was diluted in the blocking buffer according to the manufacturer’s dilution range recommendations and added to the samples for 1 hour at RT following three times washing with the permeabilization buffer and mounted in DAPI media. Images were obtained using an LSM800 Zeiss confocal microscope. RGB images captured at a magnification of at least 40X were processed for quantification of GFP-LC3 puncta per cell using ImageJ. In brief, the green channel of RGB images was extracted and converted to binary (grayscale) images by auto-thresholding, and number of GFP-LC3 granules were analyzed using the “Analyze Particles” function of the software. Colocalization analysis were performed used the ImageJ Plugin JaCoP which gives the values of two colocalization coefficients: Pearson’s correlation coefficient and the overlap coefficient ^40^.

### Proximity Ligation Assay (PLA)

The DuoLink (Merck) PLA assay was performed according to the manufacturer’s protocols. Briefly, cells were grown in 12 mm glass coverslips 24 hours before the experiment, fixed with 4% PFA for 10 minutes at RT, and permeabilized with 0,1% Triton X-100 (Sigma Aldrich) in PBS solution for 10 minutes at RT. After washing with PBS, Duolink Blocking solution was added to each coverslip and incubated in a heated humidity chamber for 1 hour at 37°C. Primary antibodies dilution was done in Duolink® Antibody Diluent (LC3B, Novus Biological NB100-2220, NF-κB p65 (F6), Santa Cruz sc-8008 and SQSTM1/p62 Enzolife BML-PW9860 diluted at 1:100) and incubated overnight at 4°C. Images were obtained using confocal microscopy (Zeiss LSM800) with 20X or 40X objectives. For quantification of immunofluorescence microscopy images, at least 30 cells were counted for each condition in each experiment. Nuclear PLA signals were measured as “corrected total cell fluorescence” (CTCF= Integrated Density – (Area of selected cell X Mean fluorescence of background readings)) using ImageJ software.

### Statistical analysis

The data obtained from three independent experiments were expressed as SEM (mean ± SEM). The analysis was performed with Prism software, version 8 (GraphPad Software, Inc). The data significance between two or more groups was calculated using unpaired t-test or one-way or two-way ANOVA. The p values were represented as statistically significant with ∗ (p ≤ 0.05), ∗∗ (p ≤ 0.01), ∗∗∗ (p ≤ 0.001), and ∗∗∗∗ (p < 0.0001).

## Data availability

All data are presented in the paper.

## Supporting information

This article contains supporting information.

## Acknowledgments

We thank Drs. Ying Zhao, Anne Simonsen and Alf Håkon Lystad for providing their materials. We are also grateful to Dr. Ivan Dikic and Dr.Grumati for sharing their protocols and materials. This work was funded in part by grants from Bundesministerium für Bildung und Forschung (BMBF) CancerSys (ProSiTu) and Helmholtz Association (iMed and SignGene) to CS.

## Author contributions

**CB**: Conceptualization, Methodology, Investigation, Data Analysis, Visualization, Project administration, Writing-Original Draft, Writing-Review & Editing. **PM** and **EK**: Methodology, Investigation, Data Analysis. **CS**: Conceptualization, Writing-Review & Editing, Supervision, Project administration, Funding acquisition.

## Abbreviations

Ang II: Angiotensin II
ATG: Autophagy-related genes
ATG16L1: Autophagy related 16 like 1
CHX: Cycloheximide
CQ: Chloroquine
CTCF: Corrected total cell fluorescence
DSS: Dextran sodium sulfate
DUBs: Deubiquitinases
EBSS: Earle’s Balanced Salt Solution
GFP: Green fluorescent protein
IBD: Inflammatory bowel disease
IEC: Intestinal epithelial cell
IKK: IκB kinase
iNOS: Nitric oxide
IR: Irradiation
IκBα: NF-kappa-B inhibitor alpha
KO: Knock-out
LAMP1: Lysosome-associated membrane glycoprotein
LC3: Microtubule-associated protein light chain 3
LIR: LC3-interacting region
LMB: Leptomycin B
LPS: Lipopolysaccharide
LSD: LIR docking site
NEM: N-ethylmaleimide
NF-κB: Nuclear factor ‘kappa-light-chain-enhancer’ of activated B-cells
NIK: NF kappa B inducing kinase
PLA: Proximity ligation assay
RelA: REL-associated protein
RHD: Rel homology domain
SQSTM1: Sequestosome 1
TLR4: Toll-like receptor 4
TNFα: Tumor necrosis factor-alpha
UBA: Ubiquitin-associated
UUO: Unilateral ureteric obstruction
WT: Wild-type

