## Supplemental information for "Autophagy limits inflammatory gene expression through targeting of nuclear p65/RelA by LC3 and p62 for lysosomal degradation"

Material included in Supplemental Information:

**S1- Autophagy induction following NF-κB activation**

**S2- p62 binds to the NF-κB p65 subunit**

**S3- p62 depletion increases early stress-induced NF-κB activation**

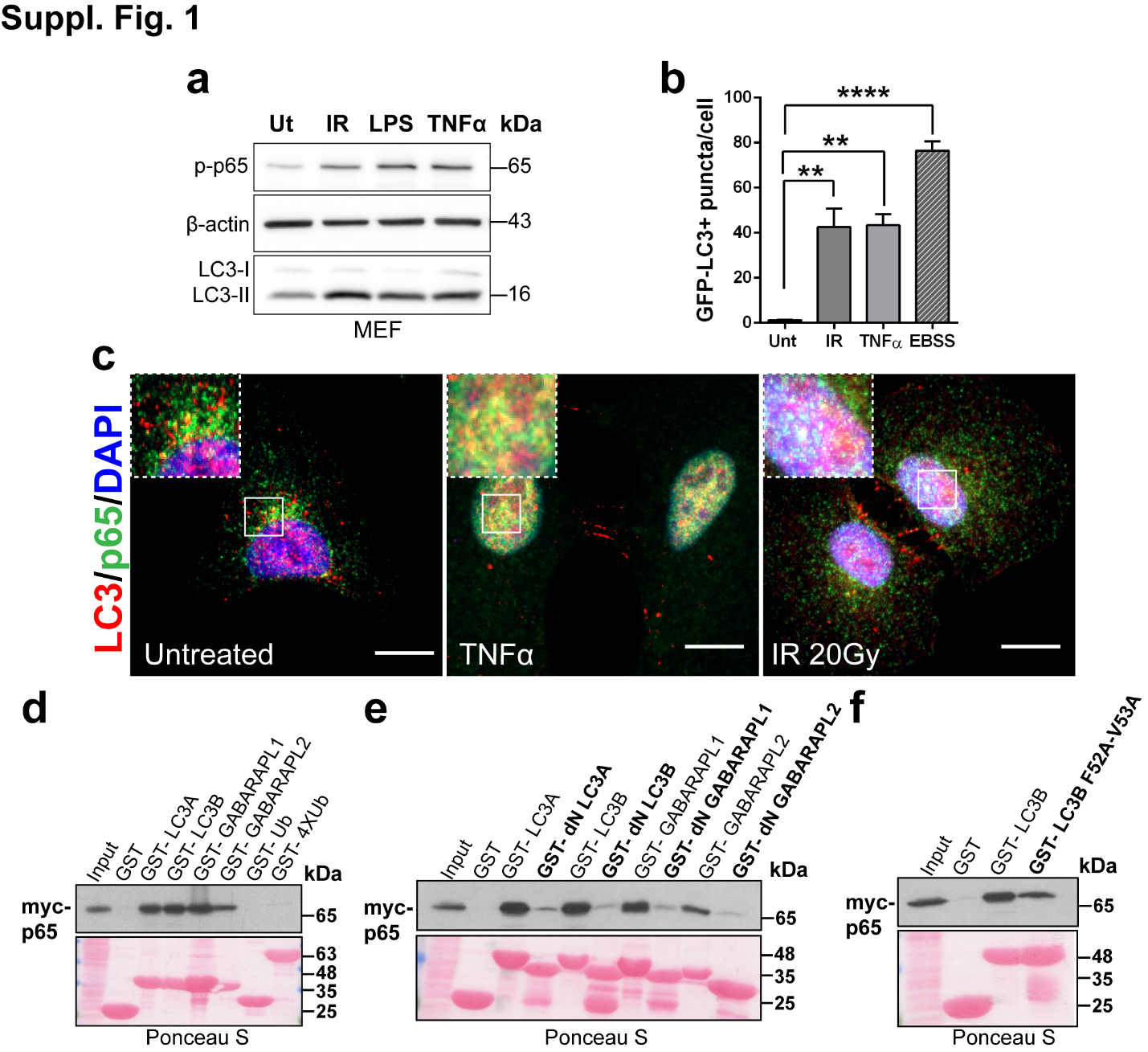

**S1. Autophagy induction following NF-κB activation: a**) Western blot (WB) analysis for the autophagy protein marker LC3 and phosphorylated NF-κB subunit p65 after 90 min irradiation (IR 80 Gy), 1 hour LPS (10 µg) or TNFα (10 ng) treatment in embryonic fibroblasts (MEF). **b**) Quantification of GFP-LC3 puncta number from the images in Fig. 1b performed using ImageJ quantification tool. Data are means ± SEM of three independent experiments with >15 cells quantified for each condition. Statistical analysis was performed by one-way ANOVA followed by Bonferroni’s multiple comparison test (**p < 0.01, ****p < 0.0001) using GraphPad Prism 8. **c**) Immunofluorescence staining for LC3 (red) and p65 (green) using A549 cells in untreated, TNFα stimulation or irradiation (IR 20Gy) conditions. Nuclei were stained with DAPI. The images shown are representative fluorescence confocal microscopic photographs of n=3 independent experiments (scale bars 10 μm). **d-f**) HEK293T cells were transfected with the myc-p65 plasmid 24 hours before analysis. Cell lysates were added to beads with immobilized GST-fusion LC3-like modifiers (GST, GST-LC3A, GST-LC3B, GST-LC3C, GST-GABARAP, GST-GABARAPL1, GST-GABARAPL2, GST-Ub, GST-4XUb (**d**) or the LC3 like modifiers lacking the unique N-terminus (dN) (e) and GSTLC3BF52A-V53A (**f**)), followed by WB using an antibody against myc- tag. Ponceau stain gels (bottom) represent the amount of GST beads constructs used for the GST-pulldown assays. All data are representative of n≥3 independent experiments.

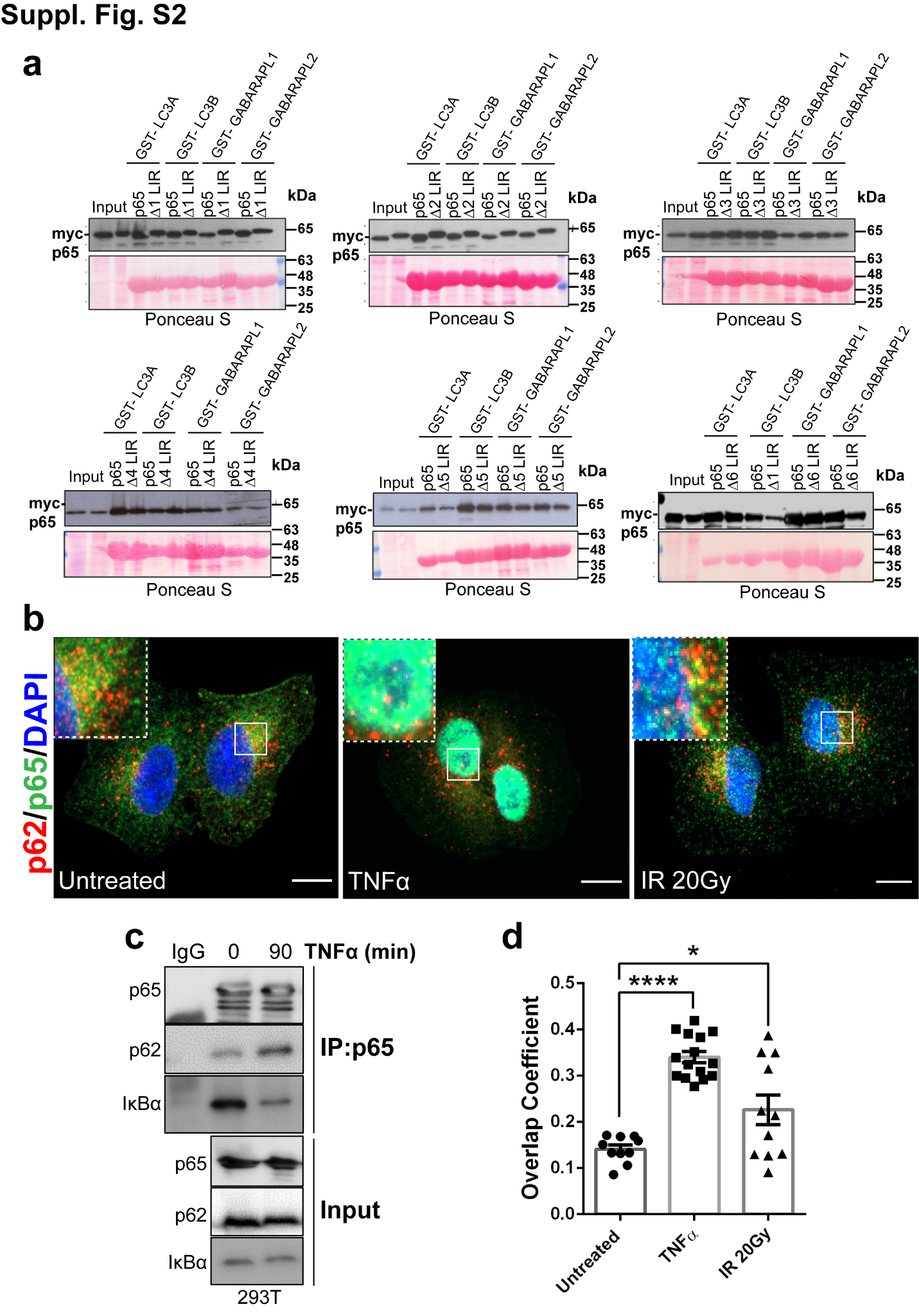

**S2. p62 binds to the NF-κB p65 subunit: a**) HEK293T cells were transfected with the myc-p65 plasmids (WT or ΔLIR mutations) 24 hours before analysis. Cell lysates were added to beads with immobilized GST-fusions of LC3-like modifiers (GST-LC3A, GST-LC3B, GST-LC3C, GST-GABARAP, GST-GABARAPL1, GST-GABARAPL2), followed by WB using an antibody against myc- tag. Ponceau stain gels (bottom) represent the amount of GST beads constructs used for the GST-pulldown assays. All data are representative of n≥3 independent experiments. **b**) Immunofluorescence staining for p62 (red) and p65 (green) using A549 cells in untreated, TNFα stimulation or irradiation (IR 20 Gy) conditions. Nuclei were stained with DAPI. Images are representative fluorescence confocal microscopic photographs of n=3 independent experiments (scale bars 10 μm). **c**) HEK293T cell extracts were harvested at 0 or 90 minutes after TNFα treatment followed by immunoprecipitation (IP) with anti-p65 antibody and WB analysis with the antibodies indicated. IκBα antibody was used as control for NF-κB activation. IgG was used as IP control.**d**) Overlap coefficient for the colocalization of PLA signals (p65:p62) with GFP-LC3 from Fig. 2e quantified using JACoP plugin of ImageJ. Data are means ± SEM of three independent experiments (n>5 cells for each condition). Statistical analysis was performed by one-way ANOVA followed by Bonferroni’s multiple comparison test (*p<0.05; ****p < 0.0001) using GraphPad Prism 8.

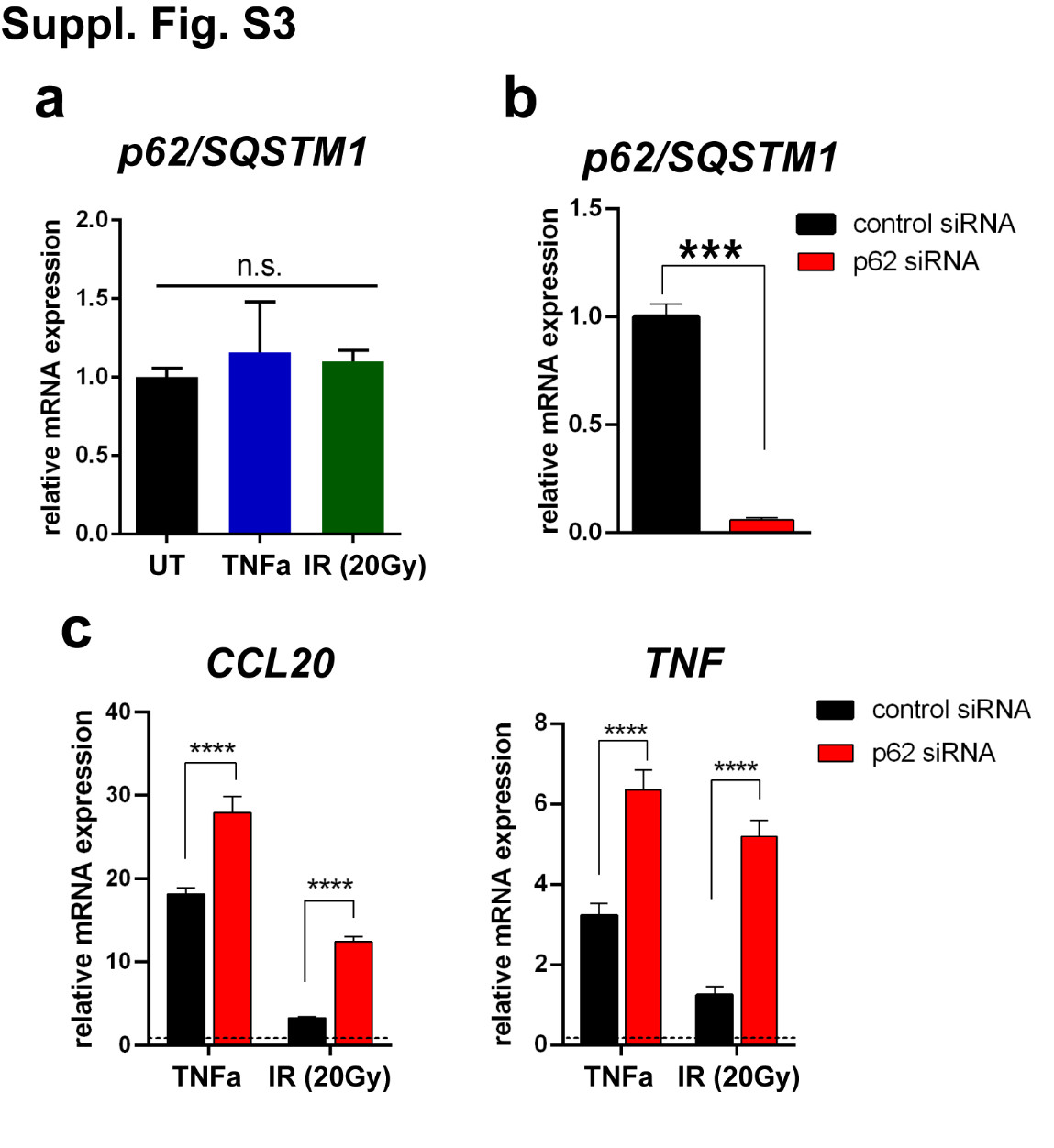

**S3. p62 depletion increases early stress-induced NF-κB activation: a**) RT-qPCR analysis of *p62/SQSTM1* expression in HEK293 cells after 5 hours of treatment with TNFα (10 ng) or irradiation (20 Gy). Results are normalized to the unstimulated (UT) control. Data are means ± SEM of three or more independent experiments. Statistical analysis was performed by one-way ANOVA followed by Bonferroni’s multiple comparison test (n.s.= not significant) using GraphPad Prism 8. **b**) *p62/SQSTM1* mRNA expression in HEK293 cells 48 hours after transfection with control siRNA or a mixture of two siRNAs targeting *p62*. Results are normalized to the siRNA control. Data are means ± SEM of three independent experiments. Statistical analyses were performed by unpaired t-test (***p < 0.001) using GraphPad Prism 8. **c**) *CCL20* and *TNFα* mRNA expression was detected by RT-qPCR in HEK293 cells expressing control or a mixture of p62/SQSTM1-specific siRNAs and analyzed 5 hours after TNFα (10 ng) treatment or irradiation (IR 20 Gy). Results are normalized to the unstimulated controls, respectively. Data are means ± SEM of three or more independent experiments. Statistical analyses were performed by two-way ANOVA followed by Bonferroni’s multiple comparison test (****p<0.0001) using GraphPad Prism 8.
